# BDNF and glucocorticoids modulate neuroplasticity via direct interaction between TRKB and glucocorticoid receptors

**DOI:** 10.64898/2026.04.08.717148

**Authors:** C.A. Brunello, M. Gil Ortiz, P. Pastor Muñoz, J.P. Araujo, J.E. Caceres Pajuelo, J.C. Avila Martí, E. Lyytikainen, S. Tonelli, G. Didio, V. Le Joncour, E. Castren

## Abstract

The overlapping effects on neuronal plasticity of acute increase in glucocorticoid levels and the BDNF-TRKB signaling indicate a deep interconnection between the two pathways. Moreover, chronic stress with elevated glucocorticoids levels and downregulation of TRKB signaling associated with reduced BDNF are both involved in the pathophysiology of different psychiatric disorders. However, the mechanism by which TRKB and glucocorticoid receptors are recruited together in the modulation of neuronal plasticity is not clear yet.

In this study we investigated the molecular mechanisms underlying the interplay of glucocorticoids and TRKB signaling *in vitro* and *in vivo*. We found that although not binding directly to TRKB, glucocorticoids promote TRKB dimerization and signaling similarly to BDNF. Moreover, the glucocorticoid receptor physically interacts with TRKB, modulating its dimerization and activity both in presence and in absence of glucocorticoids and contributing to TRKB-mediated plasticity. The transmembrane domain of TRKB is important for the interaction and for mediating the behavioral effects of TRKB and glucocorticoid receptor modulation, suggesting at least a partial overlap between the two signaling pathways. These results shed light on the interconnected effects of glucocorticoid and TRKB signaling highlighting the need for a more comprehensive understanding of the role and the dysfunction of different players contributing to synaptic plasticity.

## Introduction

Brain derived neurotrophic factor (BDNF) is a pivotal regulator of neuronal plasticity in the brain, regulating – together with the other members of the neurotrophin family – several processes including synaptogenesis and synapse consolidation, neuroprotection and survival, which eventually modulate memory, cognition and other behavioural phenotypes (Barde, 2025). In its mature form, BDNF binds to its cognate receptor tropomyosine receptor kinase B (TRKB) inducing its dimerization and cross-phosphorylation. Phosphorylated epitopes on TRKB recruit other intracellular proteins that initiate different signalling pathways that culminate in expression of genes involved in synaptic plasticity (Minichiello, 2009). Impairment in BDNF/TRKB signalling has been linked to several pathological conditions, from stress, anxiety and depression to neurodegeneration (Barde, 2025). Moreover, TRKB is the molecular target of virtually all antidepressant drugs (Brunello et al., 2024), highlighting its importance in maintaining a healthy brain. TRKB transmembrane domain possesses an inverted cholesterol recognition amino acid consensus (CRAC) domain and dimer of TRKB contains in its criss-crossed transmembrane domain a pocket for cholesterol sensing, which modulates the conformation of TRKB dimers at the plasma membrane and mediates the effects of antidepressant drugs (Cannarozzo et al., 2021; Casarotto et al., 2021). It is possible that other steroid may affect TRKB activity at the same residues.

Glucocorticoids (GC) are classical stress hormones, produced in the adrenal gland following stimuli generated through the hypothalamus-pituitary-adrenal axis. Fluctuation of GC levels are essential in maintaining a balance necessary for proper response in case of acute stress. In the brain, acute stress and temporary raising in GC level modulate synaptic plasticity and promote adaptability to novel environmental conditions and consequent lowering of GCs (Liston & Gan, 2011; Sanacora et al., 2022). On the other hand, chronic stress can lead to prolonged levels of high GC in the brain, which is a highlight of both neurodegenerative and neuropsychiatric disorders (Numakawa & Kajihara, 2023). GC exert their effects via mineralocorticoid and glucocorticoid receptors, which are classical nuclear receptor that modulate gene expression (Joëls, 2018).

Considering that BDNF and GC individually contribute to synaptic plasticity and are involved in stress-induced adaptability and resilience of neurons, it is not surprising that a growing body of evidence suggests a direct crosstalk and convergence in their signalling pathways, modulating their reciprocal effect on neuroplasticity (Duman & Monteggia, 2006; Tsimpolis et al., 2024).

Certain types of acute stress promote BDNF production (Niknazar et al., 2016; Shen et al., 2004), and acute pulses of glucocorticoids are accompanied by TRKB phosphorylation and signalling, supporting adaptive plasticity and memory consolidation (D. Y. Chen et al., 2012; Pandya et al., 2014). On the other hand, in chronic stress conditions there is an inverse correlation between GC and BDNF. Post-mortem analysis of brains of schizophrenic and depressed patients shows presence of high cortisol accompanied by low BDNF levels (Issa et al., 2010; Webster et al., 2002). In preclinical studies, GC reduced BDNF expression, indicating a direct modulation at the transcription level (H. Chen et al., 2017; Smith et al., 1995; Wosiski-Kuhn et al., 2014).

Consistently, stressed mice with impaired GR had higher levels of hippocampal BDNF compared to controls (Alboni et al., 2011). Moreover, glucocorticoids can affect cellular trafficking of BDNF: by increasing autophagy, chronic corticosterone also was shown to increase BDNF degradation and impair neurogenesis in a mouse depression model (Zhang et al., 2023). Another study showed impaired BDNF mRNA trafficking which correlated with abnormal dendritic morphology in hippocampi of chronically stressed mice (Tornese et al., 2019). Of note, in the same study ketamine, a rapid acting antidepressant, restored BDNF mRNA trafficking but not BDNF mRNA levels, indicating that localization is important for the stress response and for synaptic homeostasis. Chronic stress decreases levels and activity of TRKB (Pandya et al., 2014; Robinson et al., 2021; Wosiski-Kuhn et al., 2014). The relationship is bidirectional: it was shown that BDNF can induce phosphorylation of GR at specific phosphorylation sites, thereby modulating its downstream effects (Lambert et al., 2013; Arango-Lievano et al., 2015, 2019). This phosphorylation pattern of GR is essential for transcription of neuronal plasticity genes: low BDNF levels increase desensitization of GR and vulnerability to stress, while higher BDNF levels facilitate GR-mediated signalling as well as the response to antidepressants (Arango-Lievano et al., 2015).Additionally, exogenous BDNF can rescue the effects of chronic corticosterone both at the molecular (Zhou et al., 2000) and the behavioural level (Gourley et al., 2008).

These data indicate that coordinated glucocorticoid-BDNF signalling is essential for adaptive stress responses and for understanding the neurobiological basis of stress-related disorders. In this study, we investigate the effects of acute dexamethasone (dex) treatment, a potent glucocorticoid with high affinity for GR, on TRKB activity.

## Materials and methods

### Cell culture and treatments

Primary neuronal cultures were prepared from rat embryos at E18 as describes in literature (Sahu et al., 2019) and maintained in neurobasal supplemented with 2% B27, 1% penicillin/streptomycin and 1% L-glutamine (all from Thermo Fisher Scientific). Cell lines (Neuro2A/N2A and HEK293T) were obtained from ATCC (CVCL_0470) and cultured in Dulbeccós modified Eaglés medium (DMEM) supplemented with 10% fetal bovine serum, 1% penicillin/streptomycin and 1% L-glutamine. Both primary cultures and cell lines were maintained at 37 °C with 5% CO_2_.

Transfection of plasmids in N2A and HEK293T cells was done with Lipofectamine 2000 (Thermo Fisher) according to manufacturer’s instructions.

Dexamethasone (Sigma, #D4902), mifepristone (Sigma, #8046), BSA-Dexamethasone (Steraloids, #P0516-000) were dissolved in DMSO for *in vitro* experiments. BDNF (Peprotech, #450-02), biotin-BDNF (R&D Systems, #BT11166-025) and biotin-fluoxetine (Bosche Scientific, #H6995 biotinylated with EZ-link NHS-PEG4 Biotinylation kit, #21445, Thermo Scientific) were diluted in PBS.

#### PCA

Protein-fragment complementation assay (Remy & Michnick, 2006) was performed on N2A cells in 96 well plates as previously described (Casarotto et al., 2021; Merezhko et al., 2018). Briefly, 10.000 cells/well were transfected 24h post-plating with chimeric protein of interest (TRKB, GR, LR.FYN, PLCγ) carrying a split humanized *Gaussia princeps* luciferase at the C-terminus as a reporter protein. The PCA plasmid pairs transfected were: phGLuc(1C)-TRKB:phGLuc(2C)-TRKB; phGLuc(1C)-TRKB:phGLuc(2C)-PLCγ; PhGLuc(1C)-TRKB:phGLuc(2C)-GR; phGLuc(1C)-LR.FYN:phGLuc(2C)-TRKB; phGLuc(1C)-LR.FYN:phGluc(2C)-GR; phGLuc(1C)-TRKB(TM.TRKA):phGLuc(2C)-TRKB(TM.TRKA); phGLuc(1C)-TRKB(Y433F):phGLuc(2C)-GR.

Treatments (dex 1µM and BDNF 20 ng/ml 30 min, mifepristone 10 µM, 1h pre-treatment) were performed 48 h post-transfection in phenol-free DMEM (Lonza) and bioluminescence was measured upon addition of the substrate native coelenterazine (25 µM, Nanolight technology) with a plate reader (Varioskan Flash, Thermo Scientific). Following dimerization of two proteins of interests tagged with complimentary portions of the luciferase, the enzyme reconstitutes in its original and functional conformation producing light directly proportionally to the amount of protein of interest interacting at that moment.

### Co-immunoprecipitation and western blotting

The interaction between TRKB and GR was also assessed by co-immunoprecipitation from cell and brain lysate. Similarly, TRKB phosphorylation at tyrosine 816 was assessed from immunoprecipitated TRKB in N2A and cortical neurons. and N2A cells overexpressing TRKB and GR and primary neuronal cultures treated with vehicle or dex or pre-treated with mifepristone were lysed with NP lysis buffer (20 mM Tris-HCl, 137mM NaCl, 10% glycerol, 50mM NaF, 1% Nonidet P-40), freshly supplemented with protease (Sigma, #P2714) and phosphatase inhibitors (Sigma, #P0044). Murine brain samples were lysed with the same buffer and sonicated until the tissue was completely dissolved. Lysates were cleared by centrifugation (16.000g, 4 °C, 15 min) and added in equal amounts to Recombinant Protein G-Sepharose beads (Thermo Fisher, #101242), which were previously incubated with TRKB (R&D, #AF1494) or GR (Cell Signaling, #3660S) for 2 h at 4 °C. After overnight incubation at 4 °C, beads were pelleted, extensively washed with NP lysis buffer and protein complexes were released from beads by boiling in Laemmli buffer (5min, 95 C).

Immunoprecipitated protein complexes were loaded to 4-12% Bis-Tris gels (NuPage, Thermo Scientific) for western blotting analysis. Proteins separated by SDS-PAGE were transferred to PVDF membranes, which were blocked with 1% skimmed milk and incubated with specific antibodies (1:1000 in TBST) to detect TRKB (R&D, #AF1494), GR (Cell Signaling, #3660S), phosphoTRKB.Y816 (Cell Signaling, #4168). HRP-conjugated secondary antibodies (1:2000 in TBST, Biorad) specific for the primary antibody were used for chemiluminescence detection of the proteins with a G:BOX (Syngene) imaging system. Chemiluminescence signal was normalized against the staining of the antibody used to immunoprecipitated the sample.

### ELISAs

Surface levels of TKRB and GR were assessed by ELISA (Zheng et al., 2008). Primary cortical neurons were seeded at a density of 60.000 cells/well and grown in 96 wells ViewPlates (Perkin-Elmer) coated with 10% v/v Poly-L-Lysine (Sigma). At 14 days *in vitro*, cells were treated for 30 min, washed with ice-cold PBS and fixed with 4% PFA in PBS for 20 min. Following washes and a blocking step with 5% BSA and 5% skimmed milk, cells were incubated with the primary antibody (TRKB or GR) overnight at 4 °C in blocking buffer. Next day, following washes, the secondary HRP-conjugated antibody (Invitrogen) was added (1:5000 in blocking buffer, 1 h) and finally ECL substrate (Thermo Scientific) was added to the plate to detect chemiluminescence signal generated by HRP.

Levels of phosphorylated TRKB the pan-phosphorylation site Y706 in the murine brain were assessed with a sandwich ELISA commercial kit (Cell Signaling, #7108C1) and performed according to manufacturer’s instructions.

### Ligand binding assay

Binding assays were performed in 96 well plates as previously described (Casarotto et al., 2021; Moliner et al., 2023). Briefly, plates (Perkin-Elmer, OptiPlate 96F-HB) were pre-coated with TRKB antibody (1:500, R&D Systems, #AF1494) in carbonate buffer overnight at 4 °C and blocked with 5% Normal Goat Serum in PBS 2h at room temperature on shaker. Lysates from HEK293T overexpressing GFP-tagged wild-type TRKB (approximately 150 µg of protein per well) were incubated overnight at 4 °C. Following a quick washing step with PBS, a combination of drugs including a fixed 1 µM concentration of biotinylated fluoxetine and increasing amount of cold ligand (dex 1-100 µM or as controls fluoxetine 1-100 µM was added to the wells for 1.5h, room temperature on shaker. After three PBS washes, incubation with HRP-conjugated streptavidin (Pierce, 1:10000 in 5% NGS, 1.5 h, room temperature) and three additional PBS washes chemiluminescence was measured with ECL substrate (Thermo Fisher) and a plate reader (Varioskan Flash, Thermo Fisher).

### Colocalization analysis

Colocalization of TRKB and GR at the dendritic spine level was performed as previously described (Fred, 2019). Briefly, cortical neurons were grown on poly-L-lysine-coated coverslips, treated with dex, BDNF or vehicle and fixed with 4% PFA. Coversplips were then washed and blocked in blocking buffer (5% normal donkey serum, 1% BSA, 0.1% gelatin, 0.1% Triton X100, 0.01% Tween-20 in PBS) for one hour, before incubation with primary antibodies (1:500; TRKB, R&D, #AF1494; GR, Cell Signaling, #3660S) overnight at 4 °C. Next, coverslips were washed and incubated in Alexa Fluor-conjugated secondary antibodies (for TRKB, donkey-anti-goat 488; for GR, donkey-anti-rabbit 568) for 1h at room temperature before mounting them on coverglasses with Dako Fluorescence Mounting Media (Dako, S3023). Imaging was done with a Zeiss LSM 980 confocal, with a 63x water objective at 1024x1024 pixel resolution. At least 10 Z-stacks images were acquired per image. The colocalization analysis was performed in FIJI with Coloc2 plugin after identifying single spines as region of interest, and only those that passed the Costeśs p>0.95 were included in the analysis for the calculation of the Pearsońs correlation coefficient.

### Spinogenesis

Primary hippocampal rat neurons were seeded at a density of 50,000 cells per well in 24-well plates. On DIV21, cells were treated with mifepristone (10 µM one hour prior), dex (1 µM) and 50 ng/mL for BDNF and incubated for 24 h. The following day, the neurons were fixed with 4% PFA, washed with PBS and incubated in blocking buffer (10% donkey normal serum, 1% BSA, 0.1% gelatin, 0.1% Triton X-100, and 0.05% Tween 20 in PBS) for 1 h at room temperature.

Anti-MAP2 (Invitrogen, PA5-17646) primary antibody was incubated 1:1000 in blocking buffer without detergent overnight at 4°C under agitation. Following a round of washes with PBS, the secondary antibody (Donkey Anti-Rabbit Alexa Fluor 488, 1:1000, Invitrogen) was added for one hour. For F-actin visualization, Phalloidin Labeling Probe (AF647, #A22287, Invitrogen) was diluted 1:40 in PBS and incubated for 90 minutes. A 5-min wash with 0.1M phosphate buffer was performed before mounting. on cover glasses with Dako Fluorescence Mounting Media (#S3023, Agilent).

Images were acquired with a ZEISS LSM 980 confocal microscope and a Plan-Apochromat 63×/1.4 water objective and analysed blindly for all groups in ImageJ standardizing the phalloidin channel by adjusting pixel values to a range between 15 and 150 and generating a 5 µm x 15 µm ROI on representative portion of secondary dendrites. Spines were counted manually by two independent researchers.

### Cued fear conditioning

Animal experiments were conducted according to the international guidelines for animal experimentation and the County Administrative Board of Southern Finland (ESAVI/40845/2022). C57BL/6NTTac-Ntrk2em6006(Y433F)Tac male mice (Y433F.het) and their TRKB wild type littermates were used for the cued fear conditioning experiments performed as previously described (Karpova, N. N., et al, 2011). Briefly, a conditioned stimulus (CS: tone pulsed at 1 sec., frequency 10000Hz, total duration 30sec.) was immediately followed by an unconditioned stimulus (US: electric foot shock of 1sec. duration, 0.6 mA). On day one, mice were exposed to 5 unpredictable CS-US pairs (FC; inter-trial interval: 20 – 90sec.) in Context A (29x29x35cm, transparent Plexiglass walls and metallic grid floor, white ambient noise). On day 2 and 3, one hour before both fear extinction sessions (Ext1 & Ext 2: 12 unpredictable presentations of the CS, inter-trial interval: 20 – 60sec.; in Context B: 29x29x35cm, dark opaque Plexiglass walls, grey plexiglass floors, clean standard cage bedding under the floor and white ambient noise), mice were injected with a single intraperitoneal injection of 30 mg/kg of Mifepristone (M8046, Sigma Aldrich) or Vehicle (0.5% Tween-20, 5% DMSO, in 0.9% saline). One week after the last fear extinction, mice underwent a Spontaneous Recovery session (SR), which consisted of 4 unpredictable presentations of the CS (Context B; inter-trial interval: 20 – 60sec.), and 2 h later, a Fear Renewal (FR), again 4 unpredictable presentations of the CS (Context A; inter-trial interval: 20 – 60 s).

### Statistical analysis

The data was analyzed in Prism 10.1.1 (Graphpad) by Student’s t-test (two tails), one- and two-way ANOVA, followed by Šidák, Tukey and Dunnett’s post hoc tests. Replica numbers for cell culture experiments refers to technical samples, all experiments were replicated 2-3 times. N and specific p-values are indicated in figure legends. Unless otherwise specified, data is presented as change of vehicle.

## Results

### TRKB activity is modulated by GCs

To investigate whether acute glucocorticoid signalling affects TRKB activity, we performed protein-protein fragment complementation assays in N2A cells overexpressing TRKB (Remy & Michnick, 2006). Treatment with dexamethasone (dex), a synthetic glucocorticoid, induced TRKB homodimerization, which is the prelude of its activation, after 30 min of stimulation (Figure 1A). We then tested whether dex treatment was able to increase TRKB docking to the phospholipase C γ (PLCγ), which mediates intracellular signalling (Minichiello, 2009). Dex (1 µM) increased interaction with PLCγ, similarly as TRKB natural ligand BDNF (Figure 1B). Additionally, dex increased the localization of TRKB on the plasma membrane, hence increasing the levels of the receptor available for BDNF binding (Figure 1C). However, contrary to BDNF, which initiates a positive feedback loop to further increase production of BDNF (Barde, 2025), dex did not significantly increase the release of BDNF from cortical neurons (Figure 1D), in line with what has been previously reported *in vivo* (Jeanneteau et al., 2008).

**Figure 1.**
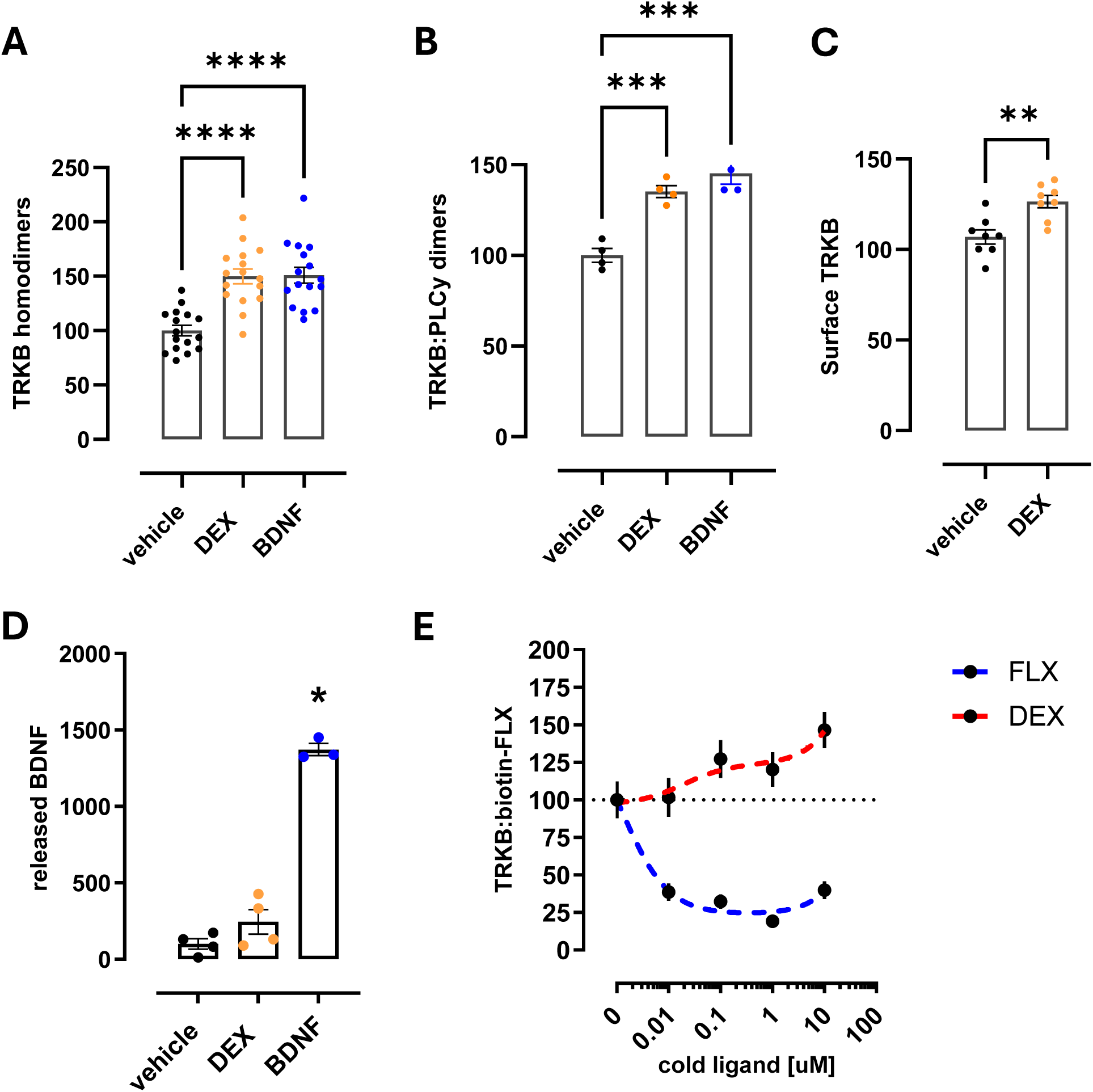
Dexamethasone modulates TRKB activity. (A) Acute 30 min administration of dex (1 μM) and BDNF (20ng/ml) induces TRKB homodimerization in N2A cells. One way ANOVA followed by Dunnett’s post-hoc test, F(3,60)=14.06 p(dex, BDNF)<0.0001. n=16/group. (B) Acute 30 min administration of dex and BDNF increases interaction of TRKB with its signaling partner PLCγ in N2A cells. One way ANOVA followed by Dunnett’s post-hoc test, F(2,9)=27.79, p(dex)=0.0007, p(BDNF)=0.0001. n=4/group. (C) Acute 30 min dex treatment increases surface localization of TRKB in hippocampal primary neurons. Unpaired t-test, p=0.002. n=8/group. (D) Acute treatment 30 min with BDNF induces BDNF release in the extracellular media as measured by ELISA (One way ANOVA, and Dunnett’s post-hoc test, p<0.0001), but dex does not significantly increases BDNF release. n=3-4/group. (E) In competition binding assay dex dose-dependently increases binding of biotin-fluoxetine to TRKB, while increasing concentration of fluoxetine displaces biotin-fluoxetine from TRKB binding. n=5.

Our previous study showed that the antidepressant fluoxetine directly interacts with the transmembrane domain of TRKB dimers and several other antidepressant drugs can displace fluoxetine from this binding site, while cholesterol increased the interaction between TRKB and fluoxetine (Casarotto et al., 2021). We performed binding assays in HEK293T cellular lysates overexpressing TRKB to assess whether treatment with increasing concentration of dex could displace biotinylated fluoxetine (Figure 1E) bound to TRKB. Dex dose-dependently increased fluoxetine binding to TRKB, while unlabelled fluoxetine disrupted binding, suggesting that dex modulates TRKB conformation in particular at the transmembrane domain, where fluoxetine has been shown to bind and act as an allosteric modulator of the receptor (Casarotto et al., 2021).

### Dex-induced modulation is mediated by GR

Since the intracellular glucocorticoid receptor (GR) is the endogenous high affinity binding partner for dex we hypothesised that the effects of dex on TRKB modulation could be mediated by the GR. Moreover, it was previously shown that TRKB physically interacts with GRs (Numakawa et al., 2009), suggesting that this interaction between TRKB and GR may be prone to modulation by glucocorticoids and BDNF. Indeed, TRKB and GR interacted in PCA in N2A cells (Figure 2A), and the interaction was increased both by dex and BDNF treatment. The interaction was confirmed on coimmunoprecipitation in the same cells (Figure 2B, 2C). On the other hand, when cells were exposed to dex conjugated to BSA and prevented dex from crossing the plasma membrane, there was no change in TRKB:GR interaction, suggesting that dex has to pass through the plasma membrane to induce its effects (Figure 2A).

**Figure 2.**
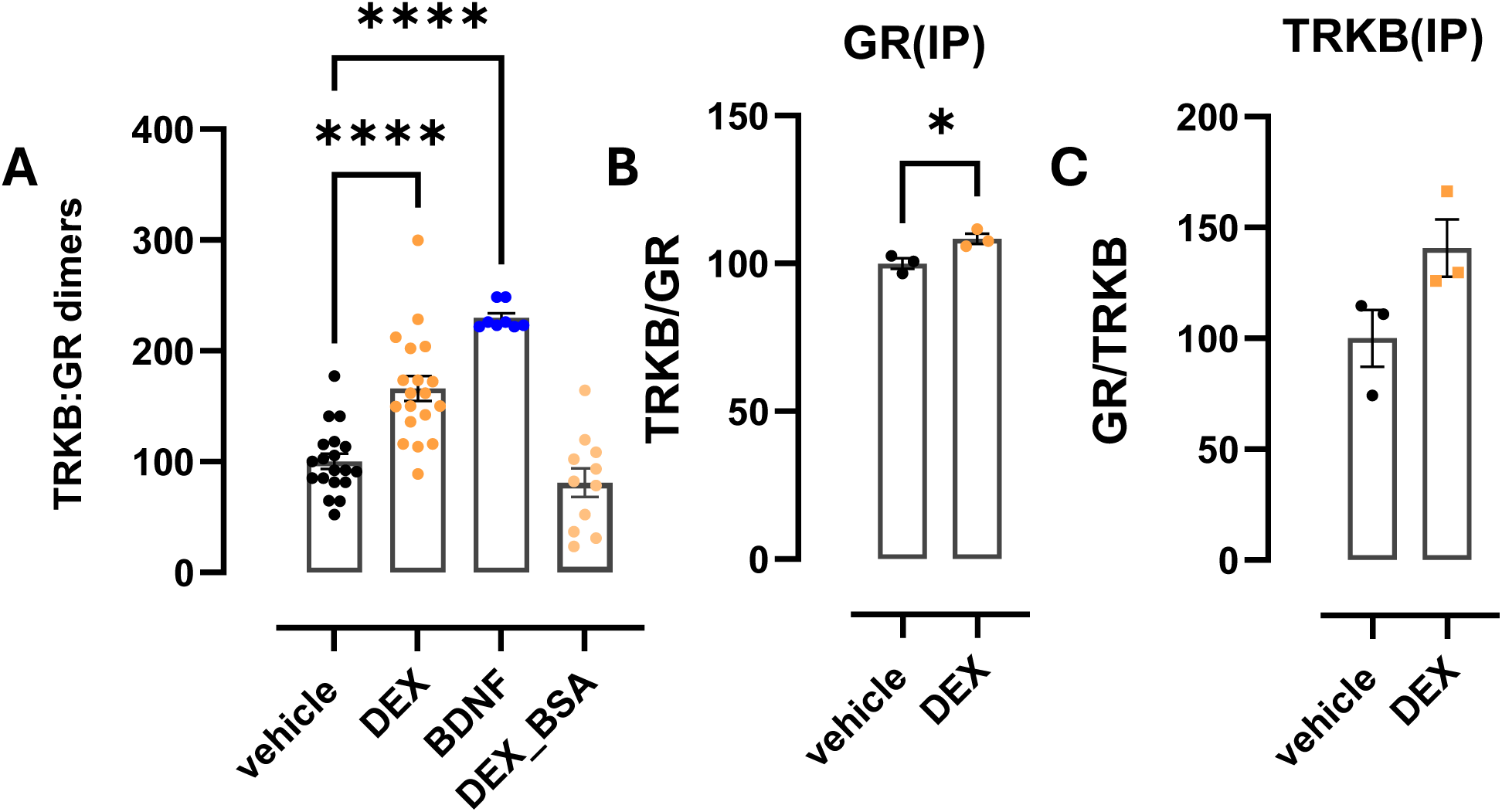
TRKB interacts with the glucocorticoid receptor. (A) Acute 30 min treatment with dex (1 μM) and BDNF (20ng/ml) increases interaction between TRKB and GR in PCA in N2A cells, while membrane impermeable dex-BSA (1 μM) does not have any effect. One way ANOVA followed by Dunnett’s post-hoc test, F(3,53)=32.61, p(dex, BDNF)<0.0001. n=8/19 per group. (B) TRKB was co-immunoprecipitated together with GR in N2A cells, and dex 30 min treatment could increase the interaction. Unpaired t-test, p=0.0272, n=3. (C) Interaction between TRKB and GR in TRKB-pulled co-immunoprecipitation, while showing a trend, was not significantly increased by dex 30 min treatment. Unpaired t-test, p=0.08, n=3.

Next, we investigated whether modulation of the activity of GRs can in turn affect TRKB activity. We pretreated N2A cells with mifepristone, the classical competitive antagonist of GR, and assessed TRKB activity with PCA. As expected, mifepristone abolished the effects of dex on TRKB dimerization (Figure 3A) indicating that functional GRs are required for dex-induced increase of TRKB dimerization. Surprisingly, mifepristone also blunted, although it did not completely abolish, the effects of BDNF on TRKB dimerization (Figure 3B). This may suggest that the interaction between GR and TRKB is also important for a correct function of TRKB in absence of glucocorticoids. Comparable effects were observed when investigating TRKB dimerization with its signalling partner PLCγ (Figure 3C). We also treated primary cortical neuronal cultures with dex or BDNF after a pre-treatment with mifepristone and checked the phosphorylation level of TRKB at tyrosine 816 with immunoprecipitation (Figure 3D). Both dex and BDNF significantly increased TRKB phosphorylation and the effects were blunted by mifepristone, although the inhibition of the BDNF effect was less pronounced than that in N2A cells. To confirm that dimerization and phosphorylation of TRKB, which are early events of synaptic plasticity, translate in structural changes at the neuronal level, we performed a spinogenesis experiment stimulated by dex and BDNF (Figure 3E, 3F). After 24 hours, BDNF and dex alone increase the spine density on primary cortical cells, as previously shown (Alonso et al., 2004; Jafari et al., 2012; Ji et al., 2005; Komatsuzaki et al., 2005), while mifepristone pretreatment inhibited the effect of both treatments on spine formation (Figure 3E, 3F).

**Figure 3.**
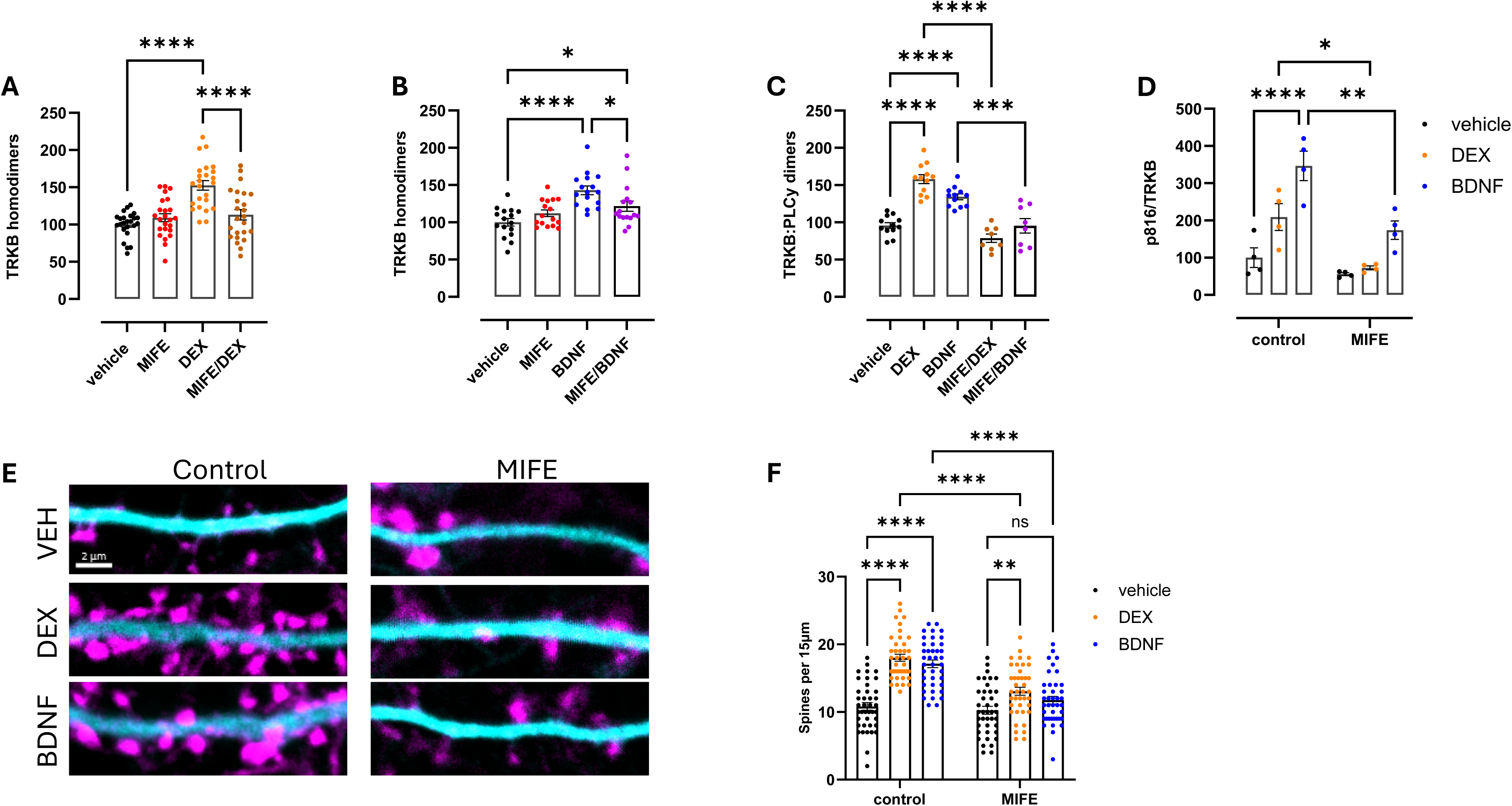
Inhibition of GR affects TRKB activity. (A) Acute 30 min treatment of dex (1 μM) increases TRKB homodimerization in PCA in N2A cells. Pretreatment with mifepristone (1 hour, 10 μM) abolishes the effects of dex on TRKB dimerization. One way ANOVA followed by Tuckey’s post-hoc test, F(3,92)=16.97, p(****)<0.0001. n=24. (B) Acute 30 min treatment with BDNF (20ng/ml) promotes TRKB homodimerization in PCA in N2A cells. Pretreatment with mifepristone (1 hour, 10μM) decreases TRKB dimerization without completely abolish it. One way ANOVA followed by Tuckey’s post-hoc test, F(3,60)=10.69; p(****)<0.0001; p(*)<0.05. n=16. (C) Similarly, as in panel A and B, PLCγ interaction with TRKB in PCA in N2A cells is abolished (dex, 1 μM) and impaired (BDNF, 20ng/ml) by mifepristone pretreatment (1 hour, 10 μM). One way ANOVA followed by Tuckey’s post-hoc test, F(4,47)=33.46 p(****)<0.0001; p(***)=0.0002. n=8. (D) In primary cortical neurons TRKB phosphorylation at Y816 induced by 30 min dex (1 μM) and BDNF (20ng/ml) is significantly reduced by mifepristone pretreatment (1 hour, 10 μM). Two way ANOVA followed by Sidák’s post-hoc test, F(2,18)=3.127, p(****)<0.0001; p(**)=0.0033; p(*)=0.027. n=4. (E) Representative images of the spinogenesis experiment. MAP2 in cyan, phalloidin in magenta. Scale bar 2 μm. (F) Pretreatment with mifepristone (1h, 10 μM) on primary cortical neurons decreases spinogenesis induced by dex and BDNF. Two way ANOVA followed by Tukey’s post-hoc test, F(2,234)=11,09, p(****)<0.0001, p(**)=0.0063.

We next investigated where the interaction between GR and TRKB occurs. In fact, GR is cytosolic typically found in complex with chaperone proteins (Backe et al., 2022). Since its ligand diffuses freely across the plasma membrane, there is no reason for the classical GR to relocate to the membrane. Furthermore, TRKB is predominantly found in intracellular vesicles and only a minority of TRKB receptors are expressed at the plasma membrane (Haapasalo et al., 2002). We checked colocalization of GR and TRKB in dendritic spines of cortical neurons, however, neither dex nor BDNF changed the colocalization of the receptors, suggesting that there is no drastic re-localization of either at the whole spine level (Figure 4A, 4B). This finding also raises the possibility that a subset of GRs may be responsible for the interaction with TRKB. The existence of membrane-bound GR responsible for the fast actions of glucocorticoids that do not require gene expression and protein synthesis is well documented (Stahn et al., 2007; Tasker et al., 2006).

**Figure 4.**
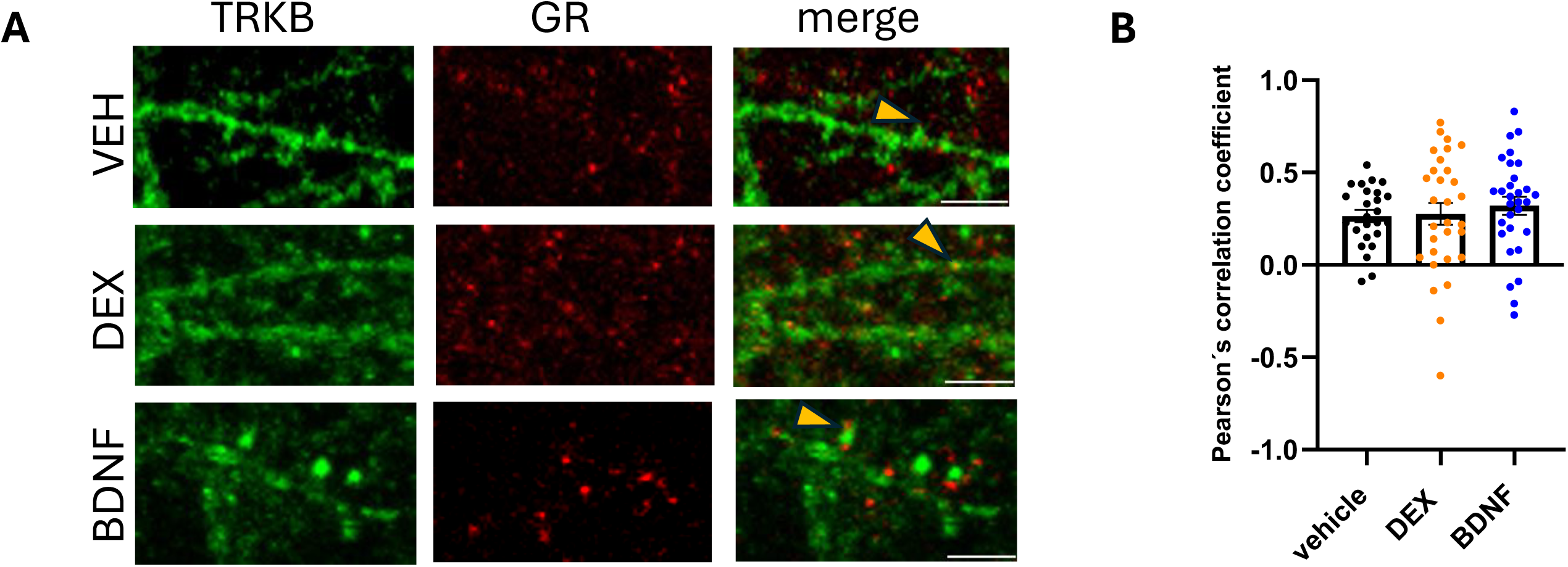
(A) Representative images of the colocalization experiment. TRKB in green, GR in red. Scale bar 4 μM. (B) Acute 30 min treatment with dex (1μM) or BDNF (20ng/ml) does not alter the colocalization of TRKB and GR in dendritic cortical spines. n=25-30 spines/group.

Although the nature of the receptor is not fully clear yet (Ping et al., 2021), it seems to share the same genomic origin (Strehl & Buttgereit, 2014) and to respond to the same markers as the classical cytosolic receptor (Rainville et al., 2019). We hypothesised that membrane GRs could be mediating the effects on TRKB.

Because dex increases binding of fluoxetine to TRKB, and fluoxetine binds to the transmembrane domain of TRKB (Casarotto et al., 2021), we hypothesised that the transmembrane domain of TRKB could be an important region for the interaction with GR as well. To investigate this, we replaced the transmembrane domain of TRKB with that of TRKA, as this domain is quite dissimilar between TRKA and TRKB. BDNF and dex fail to increase dimerization between GR and a TRKB chimera that carries the transmembrane domain of TRKA (Figure 5A). Furthermore, GR failed to interact with a TRKB receptor that carries a point mutation from tyrosine 433 to phenylalanine (Figure 5B). This tyrosine residue has been shown to be critical for the interaction of cholesterol and antidepressants with TRKB (Casarotto et al., 2021). This indicates that the transmembrane domain and its correct positioning in the plasma membrane are important for interaction with GR.

**Figure 5.**
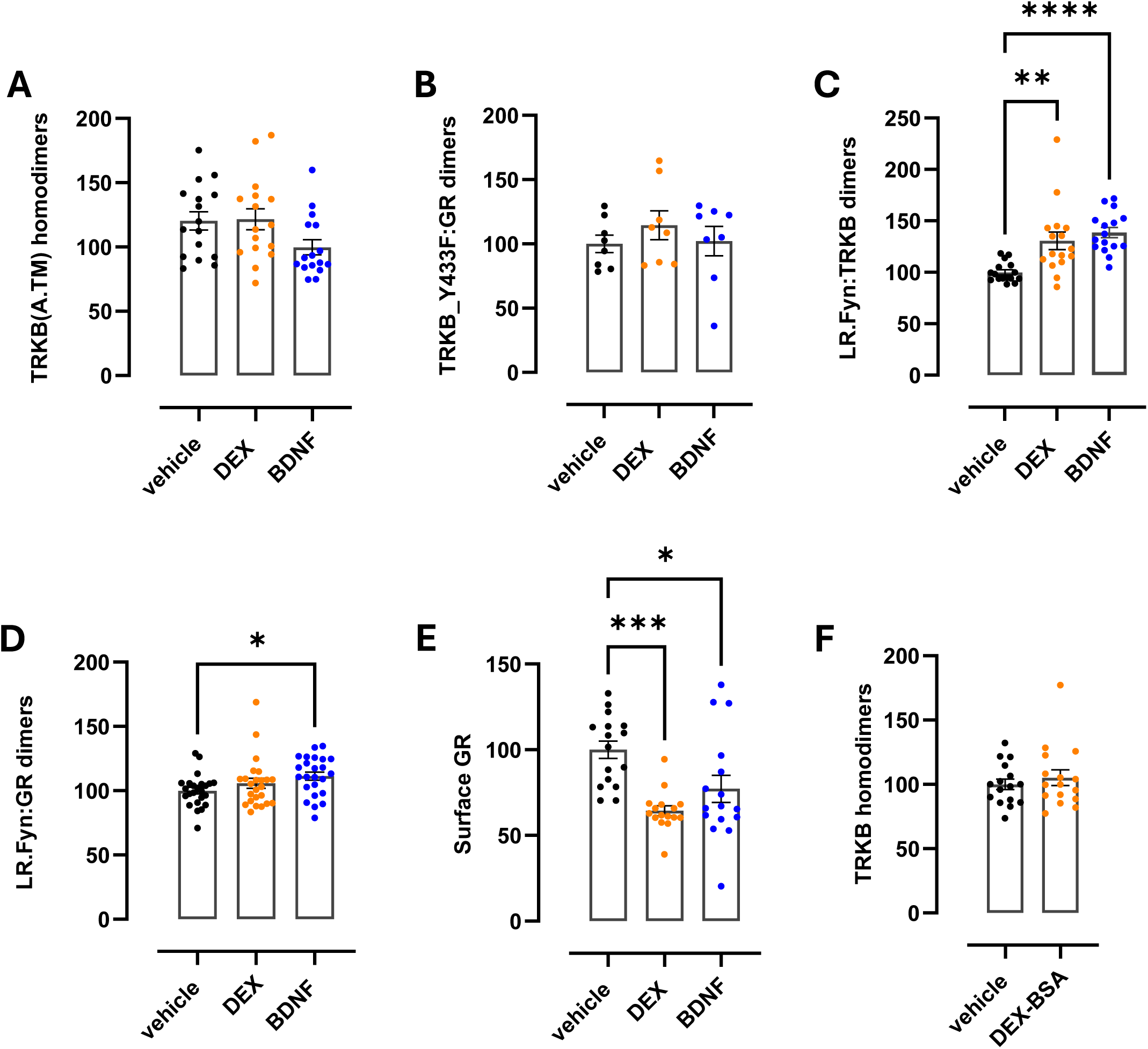
Investigation of GR:TRKB interaction. In PCA in N2A cells, acute 30 min treatment with dex (1μM) or BDNF (20ng/ml) does not induce dimerization between GR and mutant TRKB containing the point mutation Y433F in the transmembrane domain (A), or mutant TRKB that contains the whole transmembrane domain of TRKA (B). n=8/16. (C) Acute 30 min treatment with dex (1μM) and BDNF (20ng/ml) increase interaction in PCA in N2A between TRKB and a lipid raft reporter fragment from the FYN protein. One way ANOVA followed by Dunnett’s post-hoc test, F(2,45)=11.95, p(dex)=0.0012; p(BDNF)<0.0001. n=16. (D) Acute 30 min treatment with BDNF increases interaction between GR and a lipid raft reporter fragment from the FYN protein in PCA in N2A, but dex treatment does not. One way ANOVA followed by Tuckey’s post-hoc test, F(2,69)=3.095, p=0.04. n=24. (E) Acute 30 min dex (1μM) and BDNF treatments (20ng/ml) decrease the levels of membrane-exposed GR in primary cortical neurons. One way ANOVA followed by Dunnett’s post-hoc test, F(2,45)=10.16, p(dex)=0.0001; p(BDNF)=0.0124. n=16. (F) Acute 30 min treatment with membrane-impermeable dex-BSA (1μM) does not promote TRKB homodimerization in PCA in in N2A cells. n=16.

BDNF promotes translocation of TRKB to membrane-ordered microdomains called lipid rafts (Casarotto et al., 2021; Suzuki et al., 2004) that function as signalling hubs in several cellular processes (Tsui-Pierchala et al., 2002). We therefore tested whether lipid rafts could function as a dynamic meeting hub for TRKB and GR as well. The interaction in PCA between TRKB and a lipid raft-reported fragment of the FYN protein (Merezhko et al., 2020) was significantly increased following treatment with BDNF, as expected (Suzuki et al., 2004), but also after dex treatment (Figure 5C). On the other hand, the interaction between the reporter and GR was slightly increased by BDNF treatment but not by dex treatment (Figure 4D). This indicates that both dex and BDNF can mobilize the receptors to lipid rafts, but these treatments have only a slight effect on the raft localization of GR alone.

We checked the surface expression levels of GR using ELISA and found them reduced after both BDNF and dex treatments (Figure 5E), supporting the hypothesis that membrane GRs respond acutely to dex and BDNF treatments and modulate TRKB. Moreover, membrane-impermeable dex-BSA could not induce TRKB dimerization (Figure 5F). This suggests that dex permeates the cell membrane partially or completely, binds the membrane-bound GR which in turn interacts with TRKB and the two receptors are possibly internalized and recycled.

### TRKB:GR interaction could influence learning behavior

We then investigated whether the interaction between GR and TRKB, by inhibiting GRs with mifepristone, influences plasticity-dependent behaviour in a model of cued fear conditioning (Karpova et al., 2011). Because of the relevance of the transmembrane domain, we included in the study heterozygote mice for the Y433F mutation (Casarotto et al., 2021) to compare the effects of mifepristone on wild-type and mutant TRKB. Since mifepristone does not only target GRs, but also progesterone receptors (J. Chen et al., 2014), we performed the *in vivo* experiment on male mice to minimize the behavioural effects of the drugs through other steroid receptors. The rationale of the study is shown in Figure 6A. All four groups conditioned to fear by using a sound cue followed by an electric shock (Figure 6B). Mifepristone was injected acutely before the two extinction trainings, but no differences were detected among the groups in terms of freezing time, and all groups reduced freezing during the trials, indicating adequate fear extinction learning (Figure 6C). However, during the fear reinstatement test the control wild-type mice froze for a longer period of time during the first trials, an effect blunted by mifepristone, as well as in TRKB Y433F mutants, indicating that GR-TRKB interaction in the wild-type mice promotes fear reinstatement and is necessary for the retention of conditioned learning in the long-term. (Figure 6D). Mifepristone decreased the interaction between TRKB and GR in hippocampal tissue extracts from the wild-type mice, but not from the TRKB-Y433F heterozygous animals (Figure 6E), corroborating the *in vivo* results and the *in vitro* data.

**Figure 6.**
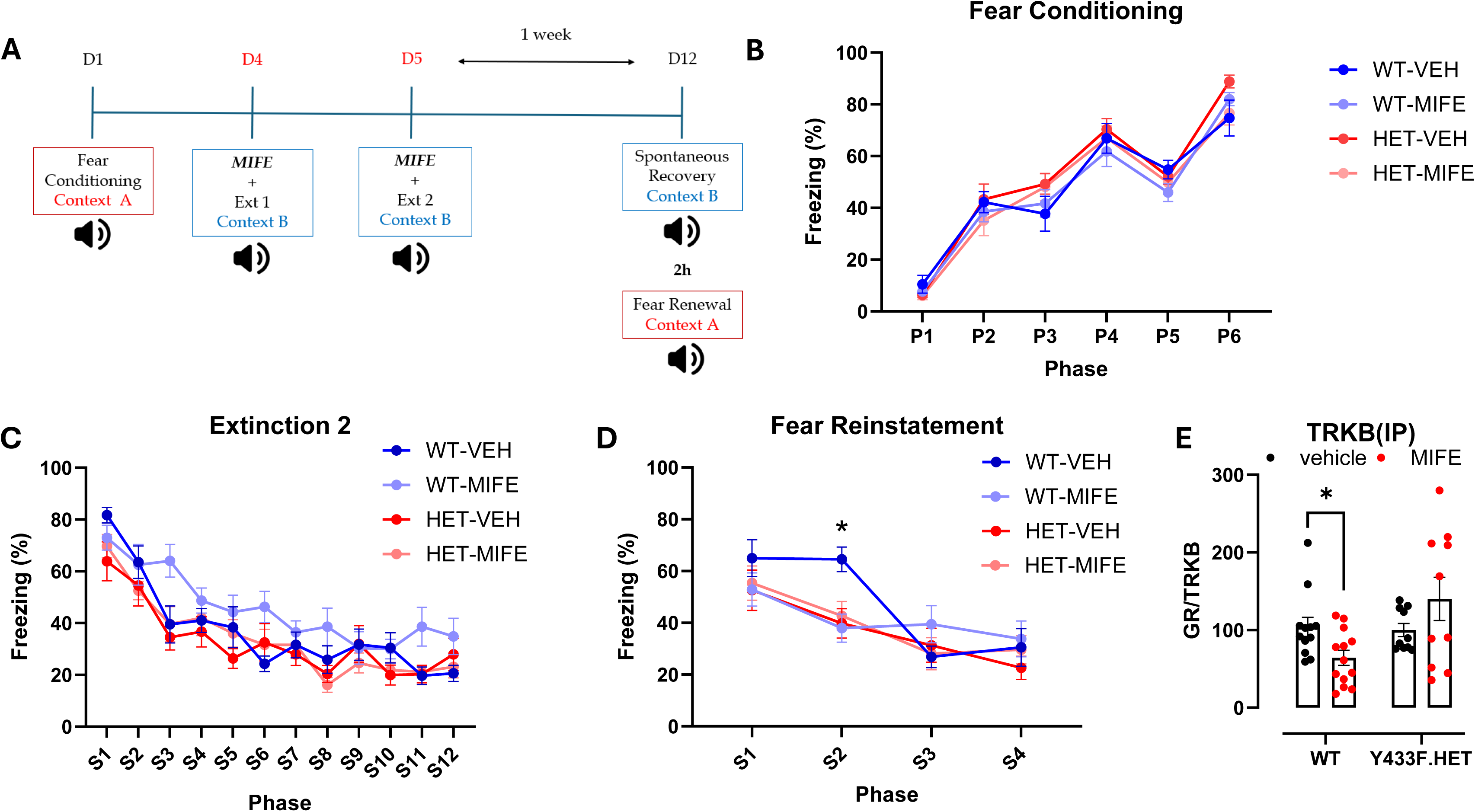
The effect of mifepristone on a cued fear conditioning paradigm. (A) The rational and timeline of the behavioral experiment. (B) Male mice were conditioned to fear with repeated foot shocks coupled with a sound cue. All groups conditioned equally, indicating that at baseline there is no behavioral difference between wild-type and Y433F.het mice. (C) Extinction training, where mice were exposed to the sound cue but not to the foot shock in a different environment (context B) compared to where they received the conditioning. All the groups similarly extinguish the fear acquired during conditioning and no effect of mifepristone treatment (30mg/kg, one hour prior) is noticeable. (D) Mice were place back in the context they received the conditioning and were exposed to the sound cue but not to the foot shock. Wild type mice that did not receive mifepristone remember better the context and freeze longer compared to the rest of the group at the beginning of the training. Two-way ANOVA repeated measures, F(9,111)=2.087, p=0.0365. n=10/11. (E) Co-immunoprecipitation of TRKB from the hippocampus of wild type mice treated with mifepristone shows a decrease interaction with GR compared to control animals, while no difference was seen for the Y433F.het group. Genotypes were analysed separately and normalized to their own control group. Unpaired t-test, p=0.0176. n=12/13.

## Discussion

This study demonstrates that acute dexamethasone treatment promotes TRKB activity via inducing a direct interaction between TRKB and the GR. This interaction was able to modulate TRKB activity in presence of dex and BDNF both *in vitro* and *in vivo*, and is mediated by the transmembrane domain of TRKB, which identifies a new level of crosstalk between the two signalling pathways. A physical interaction between the two receptors was already described over a decade ago (Numakawa et al., 2009) and it appears important for correct PLCγ signalling in presence of BDNF. The interaction can be reduced by chronic GC treatment, implicating its relevance in maintaining physiological balance in the activity of the receptors (Numakawa et al., 2009). The evidence for a physical interaction strengthens the idea that the neurobiology of BDNF and GC in the brain are closely intertwined.

BDNF and GC-systems interact at many different levels. It has been suggested that GC can regulate TRKB activity modulating the levels of immature pro-BDNF and mature BDNF via regulating the levels of tissue plasminogen activator (tPA), responsible for cleaving pro-BDNF into BDNF, and its inhibitor (Revest et al., 2014). Low glucocorticoid levels favour tPA and consequent production of mature BDNF, which results in increased ERK1/MAPK signalling. Consistently, corticosterone increases the levels of pro-BDNF over mature BDNF (Li et al., 2019), and this imbalance has been suggested to play an important role in neuropsychiatric disorders (Lu et al., 2005) . It was also shown that hippocampal GRs were acutely coupled to activation of TRKB and downstream signaling pathways during memory consolidation, suggesting that GR can rapidly enhance TRKB activity (D. Y. Chen et al., 2012). Consistently, acute exposure to corticosterone in primary cortical neurons increased TRKB protein levels, and this effect was blocked by mifepristone (Pandya et al., 2014). In astrocytes, acute one-hour administration of corticosterone upregulates BDNF mRNA, indicating that the interconnection of GR and TRKB signaling is not restricted to neuronal cells only and adding an additional layer of complexity to understand the interaction between the GC and BDNF systems (Tsimpolis et al., 2022). However, these studies do not assess a physical interaction between the two receptors, suggesting that intermediate mechanisms contribute to their crosstalk.

Supporting the idea that other intracellular players regulate the connection between GR and TRKB signaling pathways, our data indicate that in dendritic spines the colocalization of TRKB and GR is not altered by dex and BDNF treatment. However, the resolution of confocal imaging is not sufficient in detecting changes from cytosol to membrane-proximal areas. We then hypothesised that a subtype of GR could be responsible for interaction with TRKB. The membrane GR has been implicated in fast non genomic signalling of glucocorticoids (Strehl & Buttgereit, 2014), although it has also been reported to initiate nuclear translocation of GR and gene expression through binding to DNA (Jain et al., 2005; Rainville et al., 2019). The involvement of the membrane associated GR is consistent with the fact that extensive genomic expression is likely not happening during the short treatment we administered. The membrane GR has been associated to lipid rafts (Choi et al., 2017; Jain et al., 2005), and it is involved in lipid raft formation and dynamics (Van Laethem et al., 2003; Yamagata et al., 2012). Additionally, chaperone proteins associated with GR, such as Src kinase family (Sahasrabudhe et al., 2017), are also associated with lipid rafts. TRKB is predominantly expressed outside raft domains, but it can be translocated to lipid rafts, especially following BDNF stimulation (Casarotto et al., 2021; Suzuki et al., 2004). It is therefore reasonable to hypothesise that a subset of membrane microdomain could represent the meeting hub for the two receptors. We observed an interaction between TRKB and the lipid raft reporter fragment (Merezhko et al., 2020) and its increase upon stimulation, which is consistent with this hypothesis, although the interaction of GR was increased only by BDNF treatment, possibly because the majority of membrane GR may already be associated with lipid raft rather than being translocated there upon activation as for TRKB.

In our experimental condition, BSA-conjugated dex promoted neither TRKB dimerization nor interaction with GR, which suggests that even in the scenario of a direct involvement of a membrane GR, the binding does not happen from the extracellular side. This is compatible with the current knowledge of membrane GR: since glucocorticoids can freely permeate the membrane, there could be multiple binding sites that activates the same receptor (Strehl et al., 2011; Tasker et al., 2006). It has been suggested that the dex-induced phosphorylation of TRKB was dependent on genomic rather than non-genomic effects, as not only was BSA-dex unable to induce phosphorylation, but the effect was lost with blocking of transcription and translation (Jeanneteau et al., 2008). Moreover, pulse treatment with dex induced TRKB phosphorylation only with a delay, indicating that some time was required for genomic effects to occur. However, it is possible that an early, non-genomic effect modulates the interaction between TRKB and GR, while longer timepoints are needed of genomic effects to occur and further modulate TRKB activity, potentially through increased BDNF expression and release (Arango-Lievano et al., 2015).

Our *in vivo* study indicates that mifepristone partially blocks the reconsolidation of cue-conditioned fear. Similar effects have been documented before at the same dosage of mifepristone (Flavell & Lee, 2019; Pitman et al., 2011). Interestingly, the freezing behavior of mutant Y433F.het mice (which show impaired interaction between TRKB and GR), resembles that of the mifepristone-treated group, with decreased freezing throughout the fear reinstatement, and no additional effect from the mifepristone treatment, compared to the wild-type controls. This indeed suggests that the effect seen could be dependent on the interaction between GR and TRKB. Furthermore, in the hippocampi of these animals, mifepristone decreases the interaction between TRKB and GR in wild type mice but not in TRKB.Y433F mutants, supporting the importance of GR-TRKB interaction. Previous studies have shown that hippocampal infusion of BSA-conjugated corticosterone that cannot pass cellular membranes recapitulates the effect of acute stress on memory retrieval (Chauveau et al., 2010). Another study reported that the effects on memory tests of BSA-conjugated corticosterone injected in the insular cortex were counteracted by mifepristone (Roozendaal et al., 2010). These studies suggest that membrane GRs are involved in behavioral effects of glucocorticoids. However, considering the multiple molecular targets of mifepristone and the wide range of doses that have been administered even to investigate similar behavioral paradigms (Nayana et al., 2022), the characterization of the role of GR in modulating TRKB activity *in vivo* requires further investigation.

In this study, we have identified a new acute mechanism by which glucocorticoids regulate TRKB activity, via GRs (see graphical summary). Our evidence highlights the importance of the interaction between GR and TRKB transmembrane domains for proper TRKB signaling. Moreover, these findings contribute to bridge beneficial acute and detrimental chronic effects of GC on TRKB and provide a unifying mechanism linking stress physiology to neuroplasticity with implications for understanding and treating stress-related disorders.

## Acknowledgements

The authors thank Sulo Kolehmainen, Seija Lågas, Erja Huttu and the mouse behavioural phenotyping facility (MBPF, University of Helsinki) for technical assistance, and Prof S. Michnick and Dr H. Huttunen for the original GLuc plasmids. The Castréńs lab is thanked for discussions and insights. The work was supported by the Research Council of Finland (project 1333589) and the Finnish Cultural Foundation.

## Authors contributions

CAB and EC conceptualized and designed the experiments. CAB, MG, PP, JPA, JECP, EL, JCAM, ST, GD, and VLJ performed the experiments. CAB and EC wrote the manuscript.

---

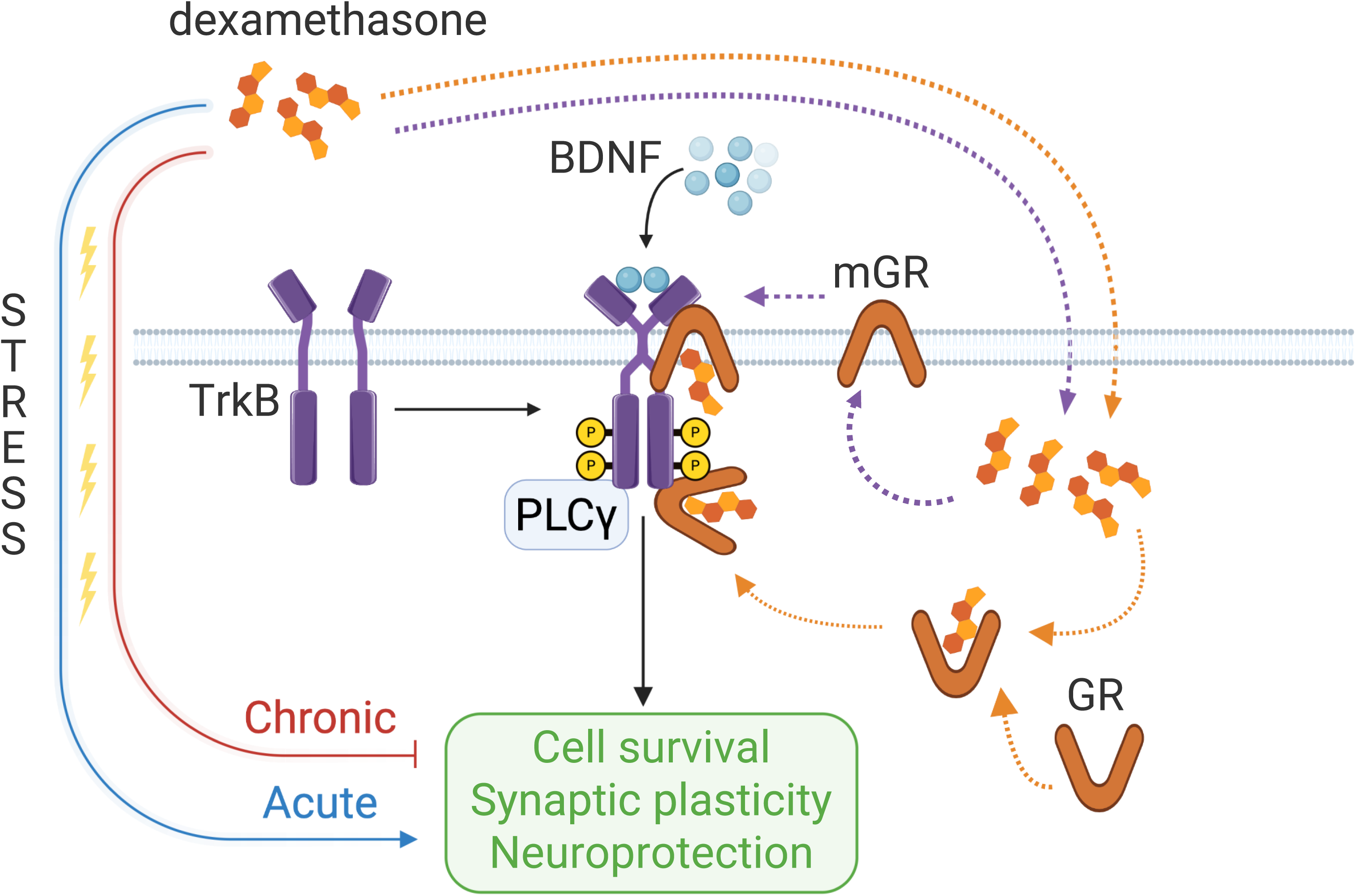
Graphical summary. Interconnection between BDNF and glucocorticoids signaling. BDNF activates TRKB, which in turn promotes neuronal plasticity and survival. Similarly, acute glucocorticoid exposure, such as dexamethasone, also induce neuronal plasticity, while chronic exposure as for instance in pathological conditions, inhibits it. Here we show that dexamethasone modulates TRKB activity by diffusing through the plasma membrane and binding the glucocorticoid receptor, possibly both the cytosolic and the membrane-bound ones. The glucocorticoid receptor interacts directly with TRKB contributing to its correct signaling.

